# A mutant bacterial O-GlcNAcase visualizes a progressive decline of protein O-GlcNAcylation in early *Drosophila* embryos critical for neurodevelopment

**DOI:** 10.1101/2022.07.04.498772

**Authors:** Yaowen Zhang, Dandan Wang, Haibin Yu, Xiaoyun Lei, Yang Meng, Na Zhang, Fang Chen, Lu Lv, Qian Pan, Hongtao Qin, Zhuohua Zhang, Daan M.F. van Aalten, Kai Yuan

## Abstract

Protein O-GlcNAcylation, a monosaccharide posttranslational modification maintained by two evolutionarily conserved enzymes, O-GlcNAc transferase (OGT) and O-GlcNAcase (OGA), is a major nutrient sensor integrating key metabolic pathways. While mutations in *OGT* have recently been associated with neurodevelopmental disorders, the dynamics and function of protein O-GlcNAcylation during early embryogenesis remain elusive. Here we develop a new fluorescent probe to visualize O-GlcNAcylation levels in live *Drosophila* early embryos. Our study shows that protein O-GlcNAcylation declines as the embryos develop to the mid-blastula transition when the facultative heterochromatin makes its first appearance. Lowering O-GlcNAcylation levels by exogenous OGA activity promotes the polycomb group O-GlcNAc protein Polyhomeotic (Ph) to form nuclear foci and K27 trimethylation of histone H3. This enhanced facultative heterochromatin formation fine-tunes the expression of several neurodevelopmental genes including *short of gastrulation* (*sog*). We provide evidence that perturbation of O-GlcNAcylation during early embryogenesis affects learning ability in adulthood, highlighting the importance of O-GlcNAcylation homeostasis for the development of the nervous system.

## Introduction

Metazoan early embryonic development is precisely choreographed to secure survival through this critical period and ensure developmental fidelity ^1^. In *Drosophila melanogaster*, embryogenesis begins with rapid, metasynchronous nuclear divisions in a single nutrient-rich cytoplasm. The first nine divisions are inside the egg and relatively rapid, after which the nuclei migrate to the surface, forming an evenly spaced monolayer of nuclei underneath the egg shell. These nuclei then divide another four times with progressively increased interphase durations (cycles 10-13, syncytial blastoderm). The completion of mitosis 13 and onset of cycle 14 mark a critical transition in development called the mid-blastula transition (MBT). During this period, the interphase is significantly extended, and surface nuclei are encapsulated by the ingression of plasma membrane to form individual cells (cellular blastoderm). Meanwhile at the transcriptome level, many maternal transcripts are degraded and zygotic transcription is widely activated. Various markers of heterochromatins are gradually emerged as individual cells start to take on distinct fates ^2,3^. The orchestra of these different early embryonic events is supported by the maternally deposited materials. Progressive degradation or dilution of key maternal gene products, such as mitotic regulators cyclins and Cdc25s, components in DNA replication machinery, as well as free histones, sends different cues to guide development ^4-6^. More recently, the consumption of maternally provided deoxyribonucleotides (dNTPs) has been reported ^7,8^ to regulate the early embryonic cell cycles, emphasizing the importance of the less understood nutrient controls on early embryogenesis.

As a major cellular nutrient sensor, protein O-GlcNAcylation is a reversible post-translational modification controlled by two evolutionarily conserved enzymes, O-GlcNAc transferase (OGT) and O-GlcNAcase (OGA). OGT catalyzes the transfer of the GlcNAc moiety from the nucleotide-sugar molecule UDP-GlcNAc to protein serine or threonine residues, whereas OGA hydrolyzes this glycosidic linkage ^9^. Given the synthesis of UDP-GlcNAc molecules via the hexosamine biosynthetic pathway (HBP) requires inputs from the metabolisms of glucose, nucleotides, amino acids, and fatty acids, the O-GlcNAcylated proteome is highly responsive to changes in nutrient availability, triggering intracellular events that regulate many biological processes ^10-12^. OGT is essential for viability of mouse embryonic stem cells and the development of mouse embryos ^13^. In the past several years, hypomorphic mutations of *OGT* have been identified in human patients with X-linked intellectual disability (XLID) ^14-17^, suggesting an indispensable function of protein O-GlcNAcylation in neurodevelopment ^18^. In *Drosophila, Ogt/super sex combs* (*sxc*) was first characterized as a member of the polycomb group (PcG) homeotic genes, and its disruption causes a super sex comb phenotype and death at the pharate stage ^19,20^. Embryos that completely lack Ogt/sxc protein arrest development at the end of embryogenesis, with homeotic transformations in the embryonic cuticle ^21,22^. To date, the dynamics of the Ogt-mediated protein O-GlcNAcylation during early embryonic development and its biological function have not been investigated. Spatiotemporal dissection of dynamic O-GlcNAcylation during embryonic development has been hindered by the lack of effective means to visualize the O-GlcNAcylated proteome in living organisms. Widely used O-GlcNAc profiling methods, such as antibody labeling, chemoenzymatic labeling, and metabolic labeling, are largely limited to imaging the O-GlcNAcylated proteome in fixed samples ^23-25^. Fluorescently conjugated wheat germ agglutinin (WGA) can be used to image glycosylated proteins, but the drawbacks are its millimolar affinity and poor specificity for the O-GlcNAc modification ^26^. Recently, a direct fluorescent labeling method for O-GlcNAcylated proteins has been developed ^27^, yet relies on metabolic assimilation of the fed fluorescent intermediates and its biocompatibility is not fully understood. Another versatile way of detecting O-GlcNAcylation is to use a mutated O-GlcNAcase from *Clostridium perfringens* (*Cp*OGA). Mutation of the catalytic residue Asp298 to Asn (D298N) inactivates the enzyme but retains its ability to bind O-GlcNAcylated peptides ^28^. Taking advantage of this property, far western and gel electrophoresis methods have been developed ^29,30^, and the mutant protein has been successfully used to enrich and profile O-GlcNAcylated proteins *in vitro* ^26^.

Here we report the development of a live imaging method for O-GlcNAcylation using the mutant bacterial *Cp*OGA. With the new approach, we depict the dynamics of the O-GlcNAcylated proteome during *Drosophila* early embryogenesis, and reveal a developmental decline of protein O-GlcNAcylation important for the timely redeployment of facultative heterochromatin. We show that forced reduction of O-GlcNAcylation level during embryonic development downregulates the expression of neuroectoderm gene *short of gastrulation* (*sog*) and affects neuronal function even in adulthood.

## Results

### A fluorescently-labeled mutant bacterial O-GlcNAcase for visualization of protein O-GlcNAcylation

To explore if the *Cp*OGA can be utilized to develop a new imaging probe for protein O-GlcNAcylation, we bacterially expressed and purified three different versions of the mutant proteins with an N-terminal HaloTag (Figure 1A). The *Cp*OGA^WT^ is a truncation mutant of the full-length OGA from *Clostridium perfringens* that can recognize and remove the O-GlcNAc moiety from protein serine or threonine residues (Figure S1A and Supplementary Table 1). The *Cp*OGA^CD^ (D298N) is catalytically dead but still able to bind to O-GlcNAcylated proteins with high affinity, and the *Cp*OGA^DM^ (D298N, D401A) loses both the catalytic and binding activities toward O-GlcNAc moiety ^26,28,29^. The purified proteins were fluorescently labeled via the N-terminal HaloTag, and the *Cp*OGA^CD^ was tested for the capability as an imaging probe using *Cp*OGA^DM^ and *Cp*OGA^WT^ as controls (Figure S1B).

**Figure 1.**
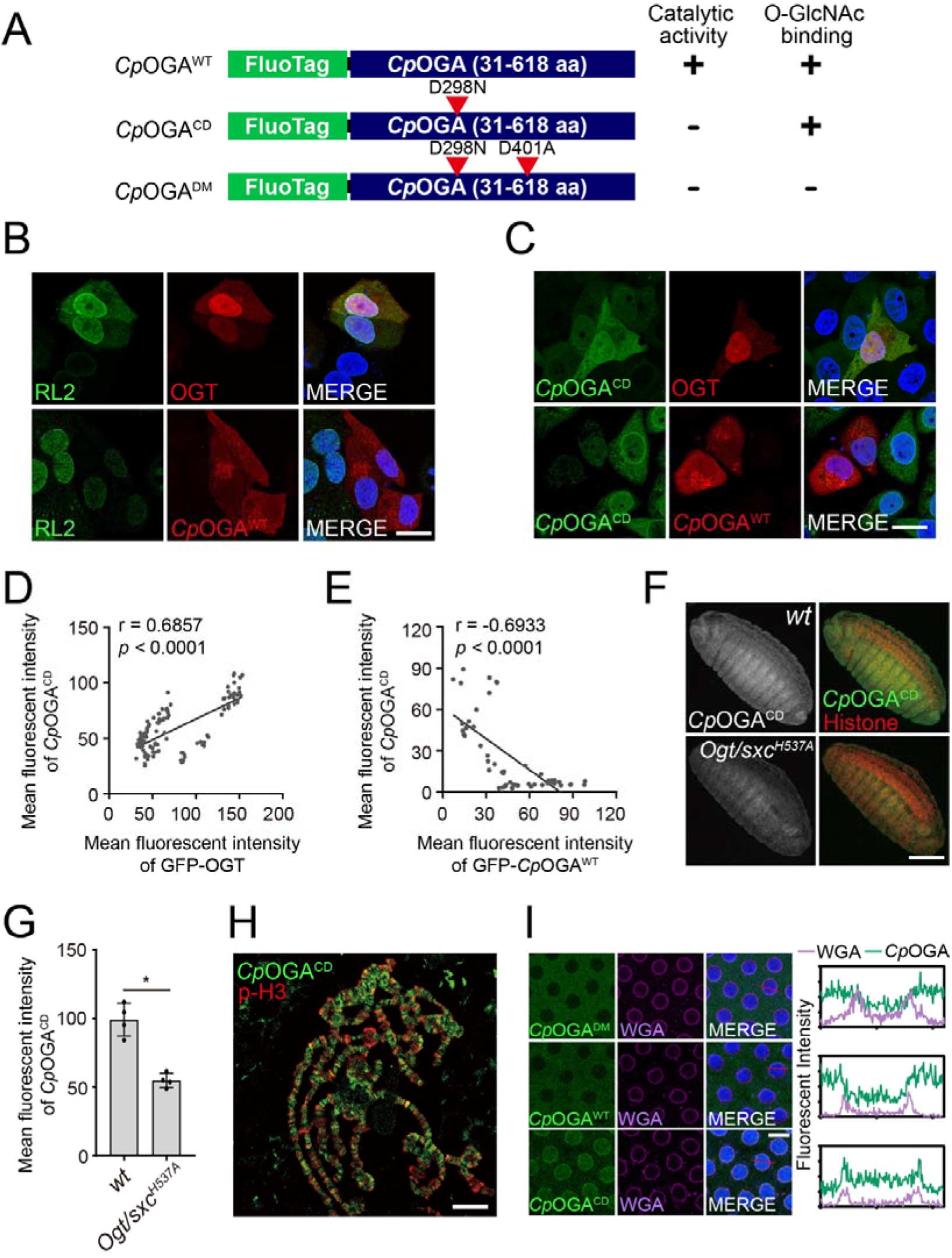
Validation of the fluorescent probe for protein O-GlcNAcylation. **(A)** Schematic of different versions of the truncated OGA from *Clostridium perfringens* (*Cp*OGA) fused to the indicated fluorescent tags. **(B-C)** Immunofluorescence of Hela cells transfected with GFP-tagged human OGT or *Cp*OGA^WT^ (red). Fluorescently labeled *Cp*OGA^CD^ protein (C) or anti-O-GlcNAc antibody RL2 (B) is used to visualize the O-GlcNAcylated proteome (green). DNA is stained with DAPI (blue). Bars: 20 μm. **(D-E)** Correlations of the fluorescent intensities between *Cp*OGA^CD^ staining and GFP-OGT or GFP-*Cp*OGA^WT^. The Pearson correlation coefficient (r) and the *p*-value are shown. **(F)** Staining of wild type (*wt*) or OGT hypomorphic mutant (*Ogt/sxc*^*H537A*^) embryos with fluorescently labeled *Cp*OGA^CD^ protein (green). Histone is shown in red. Bar: 50 μm. **(G)** Quantification of the fluorescent intensity of *Cp*OGA^CD^ staining in *wt* and *Ogt/sxc*^*H537A*^ mutant embryos. \**p* < 0.05 by unpaired t-test. Error bars represent SD. **(H)** Staining of the polytene chromosomes from the salivary gland of third instar larvae. Fluorescently labeled *Cp*OGA^CD^ protein is shown in green, and anti-phosphorylated histone H3 (p-H3) in red. Bar: 20 μm. **(I)** Live imaging of different versions of fluorescently labeled *Cp*OGA proteins (green) after injection into cycle-12 embryos. WGA is co-injected and shown in purple, and His2AvD-RFP expressed from a transgene in blue. Fluorescent intensities of WGA and the indicated *Cp*OGA along the lines (red) are plotted on the right. Bar: 10 μm.

We first tested its application in fluorescent staining of fixed samples. We transfected Hela cells with plasmids carrying human OGT or *Cp*OGA^WT^ to generate a palette of cells with varying cellular O-GlcNAcylation levels. Immunostaining with anti-O-GlcNAc antibody RL2 showed that cells transfected with OGT had stronger RL2 signals when compared with adjacent non-transfected cells. Conversely, cells that were positive for *Cp*OGA^WT^ showed reduced RL2 staining signals (Figure 1B). We repeated the experiment with the fluorescently labeled *Cp*OGA^CD^ protein (Figure 1C). The results showed that the *Cp*OGA^CD^ staining correlated positively with cellular GFP-OGT levels but negatively with that of GFP-*Cp*OGA^WT^ (Figure 1D-1E), supporting its usage as a tool to visualize the O-GlcNAcylated proteome. We noticed that the RL2 fluorescent signals were enriched in nuclei and nuclear membranes, whereas the *Cp*OGA^CD^ signals were more uniformly distributed. We also performed fluorescent staining with embryos collected from wild type and an OGT hypomorphic mutant (*Ogt/sxc*^*H537A*^) that only retained 5.6% enzymatic activity ^31^, observing stronger *Cp*OGA^CD^ signals in wild type embryos than the mutant (Figure 1F-1G). To further test the *Cp*OGA^CD^ staining on tissue samples, we prepared chromosome squash using the giant polytene chromosomes from *Drosophila* third instar larval salivary glands (Figure 1H). The *Cp*OGA^CD^ displayed a very distinct banding pattern on chromosomes as previously observed with an O-GlcNAc antibody ^20^.

We next expanded the application of *Cp*OGA^CD^ to live imaging. Wheat germ agglutinin (WGA) can bind glycoproteins and is concentrated on nuclear membranes once injected into *Drosophila* early embryos, marking O-GlcNAcylated proteins ^32^. We co-injected live *Drosophila* embryos with different *Cp*OGA mutants along with WGA, and recorded their distributions (Figure 1I). While the *Cp*OGA^WT^ and *Cp*OGA^DM^ mutant proteins showed diffuse cytoplasmic localizations, the *Cp*OGA^CD^ was enriched in nuclei and on nuclear membranes colocalizing with WGA, confirming its ability to label O-GlcNAcylated proteins in live embryos. Of note, the injection of *Cp*OGA^WT^ reduced the WGA signals on the nuclear membrane, reflecting its deglycosylation activity. The mitoses of *Drosophila* early embryos occur in metasynchronous waves initiated from the two poles ^33,34^. To assess the impact of *Cp*OGA^CD^ imaging on the progression of cell cycle and embryonic development, we injected *Cp*OGA^CD^ or WGA at one pole, and compared the interphase duration with that of the uninjected pole of the same embryo (Figure S1C). While the presence of WGA delayed local mitosis entry and disrupted the metasynchrony (upper panel, 18’00’’, region 1 versus region 2), the injected *Cp*OGA^CD^ had negligible influence on the interphase timing (Figure S1D), indicating that the *Cp*OGA^CD^ probe was better tolerated by live embryos. These results together validated the usefulness of fluorescently labeled *Cp*OGA^CD^ as a new probe for visualization of protein O-GlcNAcylation both in fixed and live samples.

### Protein O-GlcNAcylation levels decline at the mid-blastula transition in Drosophila early embryos

The maternally deposited nutrients are increasingly consumed as embryos develop, which could impact the global O-GlcNAcylation level. We injected fluorescently labeled *Cp*OGA^CD^ into developing embryos and performed live imaging to trace the developmental dynamics of the O-GlcNAcylated proteome (Figure 2A). During the syncytial blastoderm stage (C11-C13), the injected *Cp*OGA^CD^ protein was concentrated in the nuclear compartment, with prominent signals on the nuclear membrane. As the embryos enter the 14^th^ cell cycle (C14), in the first 10 minutes of interphase, the distribution of *Cp*OGA^CD^ signals remained comparable to that of the syncytial blastoderm stage. Then, the *Cp*OGA^CD^ signals, especially those in the nuclei and on the nuclear membrane, declined progressively (Figure 2B). To confirm the observed decline of protein O-GlcNAcylation around the time of MBT, we collected wild type embryos and aged them at 25 □ for 30 min, 90 min, 120 min, and 180 min respectively. The latter two time points roughly corresponded to the syncytial blastoderm and cellular blastoderm (C14) stages. Western blot with RL2 antibody or far-western blot using biotinylated AviTag-*Cp*OGA^CD^ and streptavidin both revealed marked decrease of total O-GlcNAcylation level during this developmental course (Figure 2C-2D, S2A-S2B).

**Figure 2.**
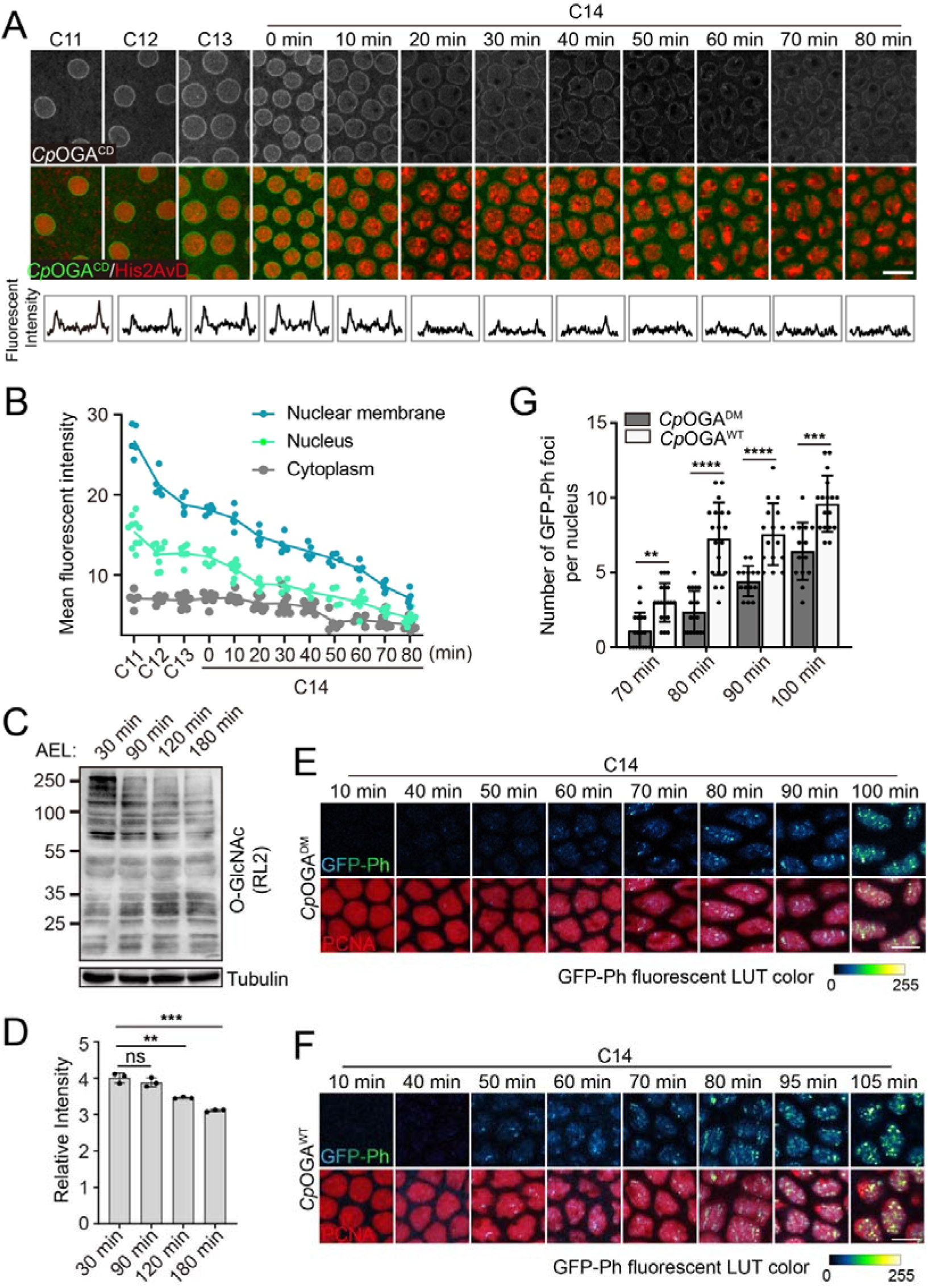
The decline in O-GlcNAcylation level and the emergence of Ph bodies in the interphase of cycle 14 (C14). **(A)** Live imaging of O-GlcNAcylation dynamics during *Drosophila* early embryonic development by injection of Alexa Fluor 488 conjugated HaloTag-*Cp*OGA^CD^ (grey/green). Nuclei are visualized by His2AvD-RFP expressed from a transgene (red). Embryonic cell cycles (C11, C12, C13) or relative times from the beginning of cycle 14 (in minutes) are shown on the top. Fluorescent intensity of *Cp*OGA^CD^ across a representative nucleus is plotted in the lower panel. Bar: 10 μm. **(B)** Quantification of *Cp*OGA^CD^ fluorescent intensity in the nucleus and cytoplasm as well as on the nuclear membrane. **(C)** Detection of total O-GlcNAcylation proteome at different timepoints after egg laying (AEL) by western blot using anti-O-GlcNAc antibody RL2. **(D)** Quantification of relative RL2 bands intensity (against tubulin). **(E-F)** Video frames showing the emergence of GFP-Ph foci in live embryos injected with catalytically active *Cp*OGA^WT^ (F) or *Cp*OGA^DM^ as a control (E). PCNA-mCherry protein is included in the injectant to visualize interphase nuclei. Relative times from the beginning of cycle 14 in minutes are given on the top. LUT: look-up table. Bars: 5 μm. **(G)** Quantification of number of GFP-Ph foci per nucleus at the indicated time points. ns: not significant, \*\**p* < 0.01, \*\*\**p* < 0.001, \*\*\*\**p* < 0.0001 by unpaired t-test. Error bars represent SD.

The *Drosophila Ogt/sxc* was originally identified as a member of the polycomb group (PcG) genes ^20,21^. One of the most well-characterized O-GlcNAcylated substrates in *Drosophila* is Polyhomeotic (Ph), the core subunit of the polycomb repressive complex 1 (PRC1) ^21,22^. Given that the PcG proteins-mediated facultative heterochromatin formation occurs in the interphase of C14 ^35,36^, coinciding with the observed decline of O-GlcNAcylation, we investigated whether the decrease in O-GlcNAcylation affected the emergence of facultative heterochromatin characteristics. During early embryogenesis, Ph progressively accumulates within polycomb bodies ^35^. We injected wild type embryos with either control *Cp*OGA^DM^ or *Cp*OGA^WT^ to deglycosylate the proteome (Figure S2C-S2D), and recorded the dynamic localizations of GFP-Ph expressed from a transgene driven by the maternal triple driver (MTD)-gal4 as previously reported ^35^. Formation of GFP-Ph foci in the nuclei was observed approximately 70 minutes after entering the interphase of cycle 14 in the *Cp*OGA^DM^ injected control embryos (Figure 2E). In the *Cp*OGA^WT^ injected embryos, however, the nuclear foci of GFP-Ph emerged much earlier (Figure 2F, 50 minutes in C14). Additionally, the number of GFP-Ph foci in each nucleus at comparable time points was also significantly increased in the presence of *Cp*OGA^WT^ protein (Figure 2G).

Ogt/sxc catalyzes O-GlcNAcylation on serine/threonine residues using UDP-GlcNAc as donor, and OGA hydrolyses O-GlcNAc from protein substrates. We injected glucosamine to boost the UDP-GlcNAc concentration and Thiamet-G to inhibit the endogenous OGA activity, so as to elevate protein O-GlcNAcylation level in the embryos. This treatment delayed the appearance of GFP-Ph foci in the interphase of cycle 14 (Figure S2E-S2F), suggesting that the assembly of Ph foci or even the repressive polycomb bodies were sensitive to the embryonic O-GlcNAcylation levels. Consistently, Polycomb (PC), another component of the PRC1 complex, also formed nuclear foci in the presence of *Cp*OGA^WT^ (Figure S2G-S2H), even though the background at this stage was high, making the visualization and quantification less accurate.

### Reduction of O-GlcNAcylation levels by exogenous OGA activity enhances facultative heterochromatin formation

The earlier emergence of Ph foci in the presence of excess deglycosylation activity suggests that there are changes in the process of facultative heterochromatin formation. To dissect this, we generated site-specific transgenic *Drosophila* stocks harboring different mutants of *Cp*OGA under the control of gal4/uas system (Figure S3A). We collected embryos expressing *Cp*OGA^WT^ or *Cp*OGA^DM^ driven by Da-gal4, and validated that the expression of *Cp*OGA^WT^ reduced total O-GlcNAcylation levels (Figure S3B). We then compared the amount of H3K27me3, a hallmark of facultative heterochromatin, between embryos expressing either *Cp*OGA^WT^ or *Cp*OGA^DM^ (Figure 3A). The *Cp*OGA^WT^ embryos had increased levels of H3K27me3 compared to the matching *Cp*OGA^DM^ embryos (Figure 3B). Western blot analysis confirmed this increase of H3K27me3 in the presence of *Cp*OGA^WT^ (Figure 3C). We also analyzed another histone modification associated with facultative heterochromatin, monoubiquitination of histone H2A at lysine 119 (H2AK119ub), however, no significant difference was observed between these two groups of embryos (Figure S3C-S3D).

**Figure 3.**
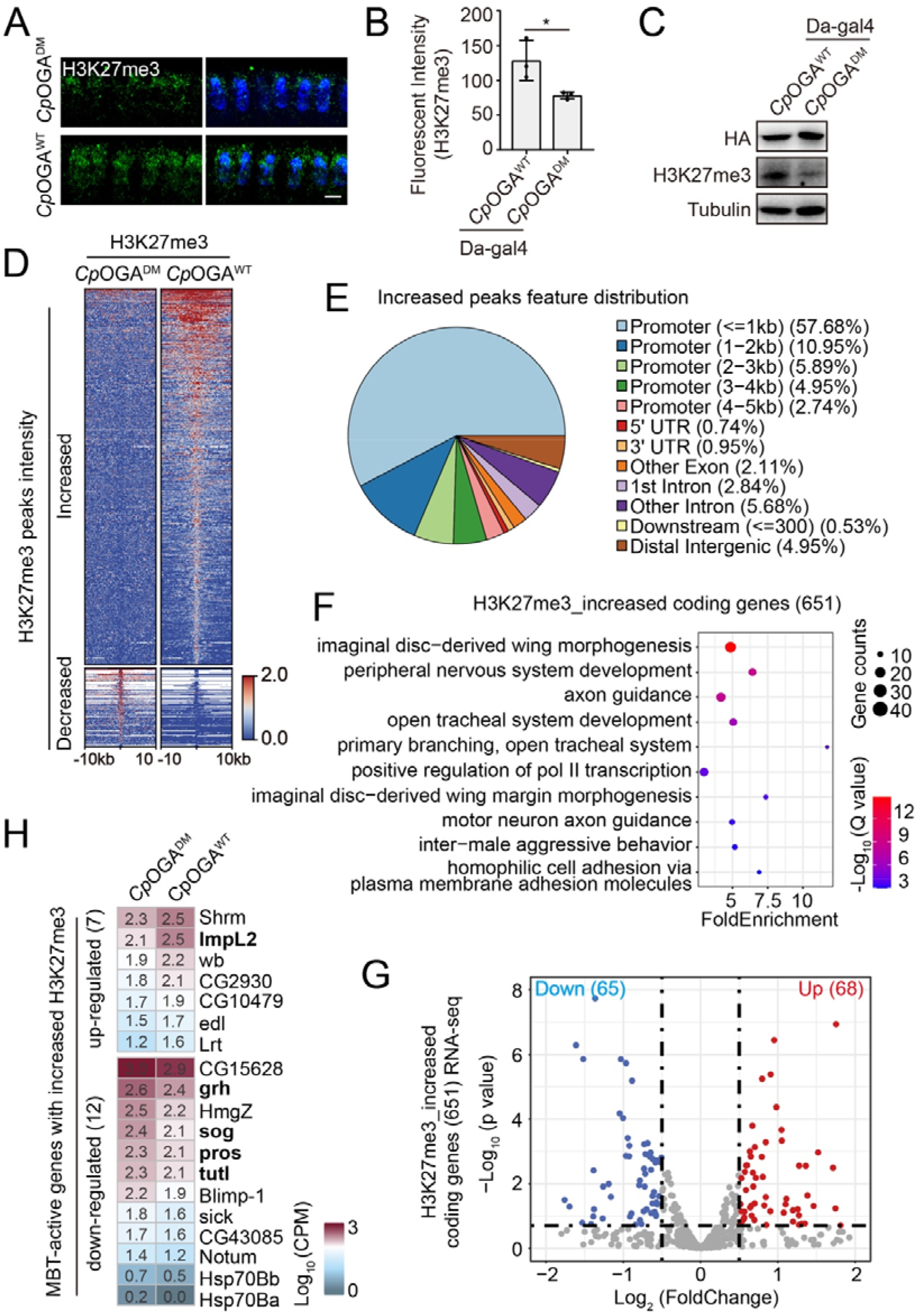
Increased H3K27me3 deposition in the presence of exogenous OGA activity. **(A)** Immunostaining of embryos expressing either *Cp*OGA^WT^ or *Cp*OGA^DM^ with anti-H3K27me3 antibody (green). DNA is visualized by DAPI staining (blue). Bar: 5 μm. **(B)** Quantification of nuclear H3K27me3 fluorescent intensity. \**p* < 0.05 by unpaired t-test. Error bars represent SD. **(C)** Detection of H3K27me3 level in embryos transgenically expressing *Cp*OGA^WT^ or *Cp*OGA^DM^. **(D)** Heatmap showing genomic regions with increased or decreased H3K27me3 ChIP-seq signals in embryos expressing *Cp*OGA^WT^ relative to that of control *Cp*OGA^DM^. **(E)** Annotation of peaks with increased H3K27me3. **(F)** Gene ontology (GO) enrichment analysis of the 651 genes with increased H3K27me3 in embryos expressing *Cp*OGA^WT^. **(G)** Volcano plot showing the differential expression of the 651 genes with increased H3K27me3. Upregulated (red) and downregulated (blue) genes are highlighted. **(H)** Heatmap representing the mRNA abundance of the MBT-active genes with increased H3K27me3 in embryos injected with *Cp*OGA^WT^ or control *Cp*OGA^DM^.

To map the increased H3K27me3 modification caused by the exogenous OGA activity, we collected embryos expressing *Cp*OGA^WT^ or *Cp*OGA^DM^ from population cages for 30 minutes and aged for 160 minutes at 25 □, and performed chromatin immunoprecipitation followed by sequencing (ChIP-seq) using anti-H3K27me3 antibody. 952 peaks were identified with increased H3K27me3 in embryos expressing *Cp*OGA^WT^ versus *Cp*OGA^DM^, and for comparison only 201 peaks showed reduced amount of H3K27me3 (Figure 3D and Supplementary Table 2). Genomic feature distribution analysis revealed that approximately 82.2% of these increased peaks were located in the promoter regions (Figure 3E). We pinpointed 651 genes according to the genomic locations of the peaks with increased H3K27me3, and found many of the genes were involved in developmental processes including wing morphogenesis as well as nervous system development (Figure 3F and Supplementary Table 3).

The facultative heterochromatin selectively silences gene expression as cells differentiate. To analyze the impact of the increased H3K27me3 deposition on the process of zygotic genome activation at this developmental stage, we carried out single embryo RNA-seq on embryos injected with either *Cp*OGA^WT^ or *Cp*OGA^DM^. *Cp*OGA proteins were injected in cycle 12 and the embryos were collected for RNA-seq approximately 1 hour after entering cycle 14 (Figure S3E-S3F). Of the 651 genes that showed increase H3K27me3 in embryos with excess *Cp*OGA^WT^, we were able to identify 65 genes that were downregulated at the mRNA level even in the presence of many confounding variables at this embryonic stage, such as degradation of maternal mRNAs as well as activation of zygotic transcription (Figure 3G and Supplementary Table 4). We selected a panel of representative genes and validated their decreased mRNA abundance using qPCR with embryos transgenically expressing *Cp*OGA^WT^ or control *Cp*OGA^DM^ (Figure S3G).

There is massive *de novo* recruitment of RNA polymerase II (Pol II) during the MBT, and a previous study has identified 251 “MBT-active” zygotic genes that are bound by Pol II and expressed at significant levels in cycle 14, as well as 593 “MBT-poised” genes that are activated later in development ^37^. Given the impact of H3K27me3 on the dynamics of Pol II ^38^, it is possible that the transcription of the 251 “MBT-active” genes is particularly sensitive to changes in H3K27me3 deposition. We found 75 of the 251 “MBT-active” genes had increased H3K27me3 in embryos expressing *Cp*OGA^WT^ (Figure S3H), and twelve of them, including *sog, pros*, and *grh*, displayed significant decreases in their mRNA levels (Figure 3H, S3I, and Supplementary Table 5).

### Embryonic O-GlcNAcylation dynamics regulates sog-Dpp signaling and is important for normal neuronal function

We turned our attention to *sog* because its gene product is a component of the evolutionarily conserved machinery for dorsal-ventral patterning ^39^, and sog opposes Dpp activity to specify the development of neuroectoderm ^40^. In embryos expressing *Cp*OGA^WT^, H3K27me3 was increased both in the gene body of *sog* and at two known regulatory loci, the distal enhancer and the intronic enhancer (Figure 4A). To test if the downregulated mRNA level of *sog* was due to impairment in transcription, we enlisted the MS2-MCP system for the visualization of nascent transcripts. Tandem MS2 hairpin sequences were placed under the control of one of the two *sog* enhancers, and the transcription driven by the enhancer recruited MCP-GFP, forming nuclear foci to report the presence of nascent transcripts (Figure 4B). In the control *Cp*OGA^DM^ injected embryos, MCP-GFP nuclear foci were visible in the interphase of cycle 13 (C13), disassembled during mitosis, and then reappeared at the beginning of cycle 14 (C14) (Figure 4C). In embryos injected with *Cp*OGA^WT^ however, the MCP-GFP nuclear foci were no longer observed in C14, indicating a premature shutdown of *sog* transcription in the presence of excess OGA activity (Figure 4D). These results consolidated the conclusion that *sog* expression was sensitive to cellular O-GlcNAcylation levels.

**Figure 4.**
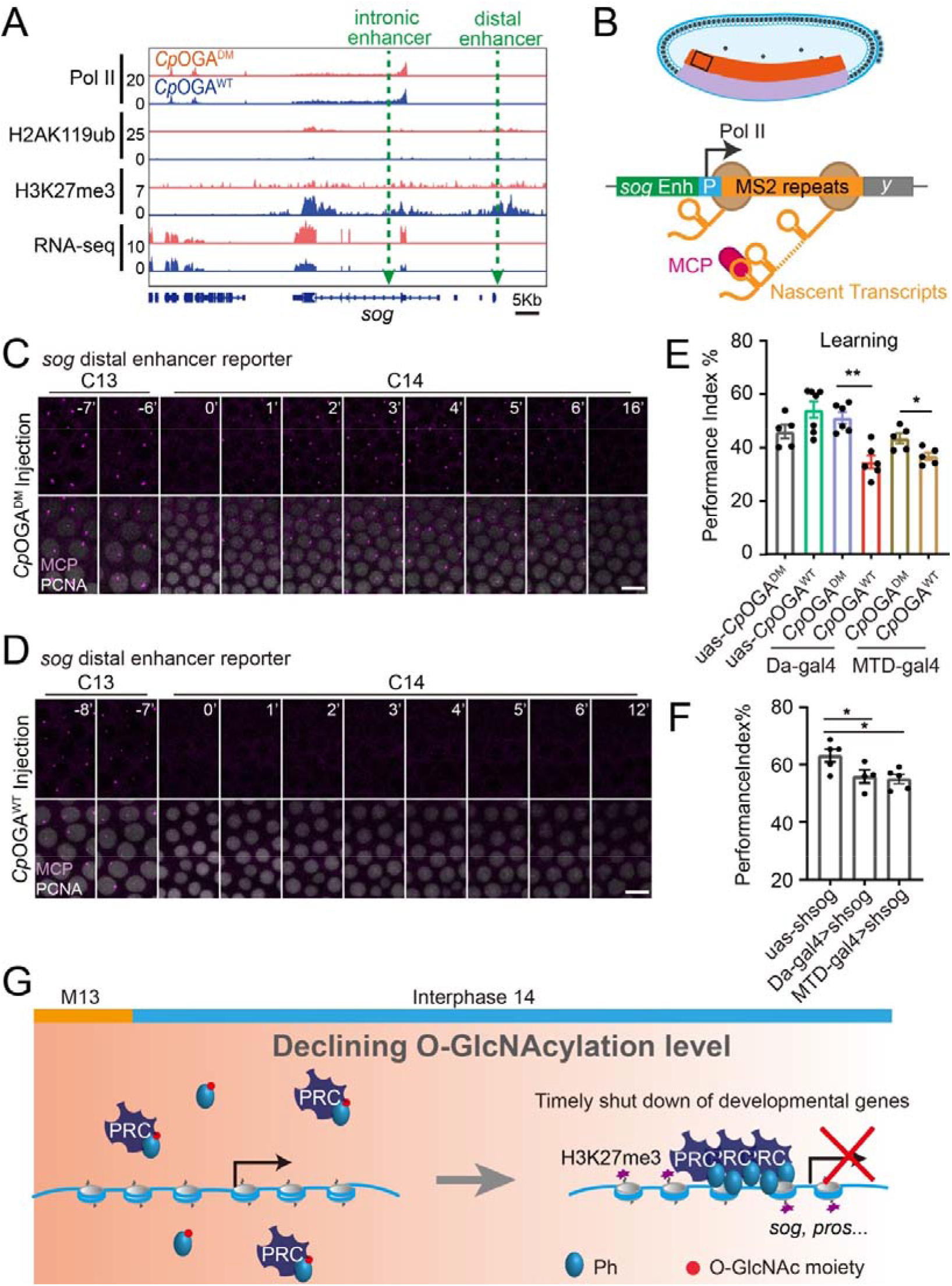
Inhibition of *sog* transcription by reducing O-GlcNAcylation levels. **(A)** Genomic snapshot showing H3K27me3, H2AK119ub, and RNA pol II ChIP-seq peaks and RNA-seq signals at the *sog* locus in embryos containing *Cp*OGA^WT^ (blue) or *Cp*OGA^DM^ (red). The two enhancers (distal and intronic) controlling the *sog* expression are labeled by green arrows. **(B)** The MS2-MCP reporter controlled by the enhancers of *sog*. **(C-D)** Video frames from live imaging of the MS2-MCP system reflecting the nascent transcription of *sog* in the presence of either *Cp*OGA^WT^ (D) or control *Cp*OGA^DM^ (C). MCP-GFP foci representing the *sog* nascent transcription are shown in purple, and PCNA-mCherry protein mixed in the injectant to visualize nuclei are in grey. Bars: 10 μm. **(E)** A compilation of performance index of the indicated flies in the learning test. **(F)** A compilation of performance index of the control and *sog* knockdown flies in the learning test. \**p* < 0.05, \*\**p* < 0.01 by unpaired t-test. Error bars represent SD. **(G)** A model summarizing the declining O-GlcNAcylation level in early embryonic development influences the formation of facultative heterochromatin and the transcription of several neurodevelopmental genes.

OGT hypomorphic mutations have been associated with X-linked intellectual disability (XLID) ^18^, suggesting that the attenuated O-GlcNAc proteome could impact normal neuronal function. Given that the excess deglycosylation activity provided by *Cp*OGA^WT^ could effectively downregulate protein O-GlcNAcylation levels, we assessed the learning ability of flies expressing either *Cp*OGA^WT^ or control *Cp*OGA^DM^ using a classical conditioning paradigm ^41^. Flies were trained to associate a given odor with electric shock once and immediately tested for odor preference in a T-maze apparatus (Figure S4C and Supplementary Movie 1). We first tested the flies expressing *Cp*OGA^WT^ or *Cp*OGA^DM^ ubiquitously driven by Da-gal4 (Figure S4A), and we observed that *Cp*OGA^WT^ expressing flies showed marked deficits in learning, confirming the importance of proper O-GlcNAcylation levels for the nervous system (Figure 4E). We next asked whether perturbations during early embryonic development influenced the neuronal function of adult flies. To this end, we drove *Cp*OGA expression with MTD-gal4 which was expressed during oogenesis and in the early embryos (Figure S4B, S4D). The expression of *Cp*OGA^WT^ induced by MTD-gal4 downregulated *sog* mRNA level (Figure S4E), and more intriguingly, these flies also manifested impaired learning ability when compared with those expressing the control *Cp*OGA^DM^ (Figure 4E). Expression of *Cp*OGA^WT^ downregulated the transcription of *sog*, which is critical for early embryonic neuroectoderm development. To test if the learning deficit induced by *Cp*OGA^WT^ was linked to the transcriptional downregulation of *sog*, we knocked down the expression of *sog* ubiquitously or only during early embryogenesis, and we observed a similar impairment in learning (Figure 4F), suggesting that the forced reduction of O-GlcNAcylation level could affect adult neuronal function via downregulating *sog* transcription during early embryogenesis.

## Discussion

Early embryogenesis is perhaps the most vulnerable period in the life cycle of a living organism. Comparing to the well-studied cell cycle remodeling and the reprogramming in transcription and epigenetics ^3,42^, we have limited understandings on the nutrient control of this developmental stage. In this report, we repurposed a mutant bacterial O-GlcNAcase, *Cp*OGA, as a tool for imaging of O-GlcNAcylation both in fixed samples and living embryos. This new imaging approach labeled the O-GlcNAcylated proteome with relatively low toxicity, enabling more complex live analysis paradigms in future O-GlcNAcylation studies. We uncovered a progressive decline in protein O-GlcNAcylation at the MBT, and observed that decreasing O-GlcNAcylation levels accompanied the emergence of hallmarks of facultative heterochromatin. Attenuation of O-GlcNAcylation levels by excess deglycosylation activity enhanced the deployment of facultative heterochromatin, inducing earlier shutdown of transcription of *sog*, a gene that specifies the development of neuroectoderm (Figure 4G). Lastly, we propose that maintaining a delicate balance of O-GlcNAcylation levels during early embryonic development is crucial for the development of nervous system, as perturbation of O-GlcNAcylation at the embryonic stage translated into learning deficits even in adulthood.

As part of the nutrient control of early embryogenesis, our results suggest a connection between protein O-GlcNAcylation and the formation of facultative heterochromatin. The first evidence of such a mechanism came from the identification of *Drosophila Ogt* as a PcG homeotic gene *super sex comb* (*sxc*) ^19^. *Drosophila* embryos lacking zygotic expression of *Ogt/sxc* died at late pupal stage, and embryos lacking both maternal and zygotic *Ogt/sxc* died even earlier at the end of embryogenesis, both with homeotic transformation phenotypes similar to that of the *Ph* mutant. Ogt/sxc was able to O-GlcNAcylate Ph on the serine/threonine-rich stretch upstream the SAM domain, and these sugar modifications prevented Ph aggregation *in vitro* ^22^. The SAM domain of Ph forms a helical multimer ^43^, and this multimerization ability was required for polycomb repression ^22^. Given that Ph was incrementally assembled into polycomb bodies in the interphase of cycle 14 ^35^, and this process was sensitive to the embryonic O-GlcNAcylation levels (Figure 4G), we propose that during early embryonic development, the progressive decline of O-GlcNAcylation allows slow and controlled polymerization of Ph to ensure faithful establishment of polycomb repression and facultative heterochromatin. Changes in the kinetics of the decline compromise the accuracy of this epigenetic control, and in extremis acute loss of O-GlcNAc modification results in disorganized polymerization and aggregation of Ph ^22^.

Our study also emphasizes the regulation of the embryonic sog-Dpp signaling by protein O-GlcNAcylation. O-GlcNAcylation seems to counter the Dpp/BMP signaling pathway at multiple layers. The UDP-N-acetylglucosamine pyrophosphorylase Mummy, a key enzyme catalyzes the formation of UDP-GlcNAc, antagonizes the Dpp/BMP signaling ^44^. Furthermore O-GlcNAcylation inhibits the Dpp/BMP type I receptor Sax ^45^. Our results reveal that at transcriptional level, low O-GlcNAcylation via polycomb repression inhibits the expression of the Dpp/BMP antagonist *sog*. Additionally, both Dpp and sog are modified by N-linked glycosylation to control their activities in the extracellular milieu ^46-48^. Taken together, these data suggest a relationship between glycosylation or to some extent the nutritional status and the Dpp/BMP signaling pathway. The evolutionary origin and molecular details of this relationship are worthy of future investigation.

OGT is a multi-domain protein, with an N-terminal tetratricopeptide repeat (TPR) domain that mediates interactions with binding partners and substrates, and a C-terminal catalytic domain harboring the O-GlcNAc transferase activity. Mutations in both the TPR and catalytic domain have been identified in patients with XLID ^14-17^. In cultured patient cells, global O-GlcNAcylation level was nearly unaltered, likely being compensated by downregulation of OGA. It is therefore not yet clear whether changes in the O-GlcNAcylated proteome underpin the XLID phenotypes. Our results using *Drosophila* models ectopically expressing *Cp*OGA^WT^ show that lowering O-GlcNAcylation level is sufficient to cause learning deficits, and moreover, lowering O-GlcNAcylation level during early embryonic development can impact learning in adult flies. Our findings cast new light on the etiology of the *OGT* associated XLID, and suggest that the misregulated Dpp/BMP signaling pathway could be involved in the neurodevelopmental defects.

## Acknowledgements

We gratefully acknowledge Drs. Hong Xu, Zongzhao Zhai, Giacomo Cavalli, Liming Wang, Michael Levine, Hernan Garcia, the Developmental Studies Hybridoma Bank, the Bloomington *Drosophila* Stock Center, the core facility of *Drosophila* resource and technology at SIBCB, and TsingHua Fly Center for antibodies and fly stocks. We thank colleagues in the center for medical genetics, members of the Yuan lab, and Ignacy Czajweski for discussions. This project has been supported by the National Natural Science Foundation of China (grants 91853108, 92153301, 31771589, and 32170821 to K.Y, 32101034 to F.C), Department of Science & Technology of Hunan Province (grants 2017RS3013, 2017XK2011, 2018DK2015, 2019SK1012, and 2021JJ10054 to K.Y, and the innovative team program 2019RS1010), and Central South University (2018CX032 to K.Y, 2019zzts046 to Y.Z, 2019zzts339 to X.L, 2021zzts497 to H.Y, and the innovation-driven team project 2020CX016). K.Y is supported by the National Thousand Talents Program for Young Outstanding Scientists. D.M.F.v.A. is supported by the Hunan provincial Hundred Talents Program for Foreign Experts.

## Author contributions

Conceptualization: D.M.F.v.A., K.Y.; Methodology: Y.Z., H.Y., F.C., H.Q., K.Y.; Validation: Y.Z., D.W., H.Y., Y.M., F.C., L.L.; Software: Y.Z., X.L.; Formal Analysis: Y.Z., H.Y., X.L., F.C., K.Y.; Investigation: Y.Z., D.W., H.Y., Y.M., N.Z.; Resources: Q.P., Z.Z., H.Q., K.Y.; Data Curation: Y.Z., D.M.F.v.A., K.Y.; Writing-Original Draft: Y.Z., K.Y.; Writing-Review & Editing: D.M.F.v.A., K.Y.; Visualization: Y.Z., D.W., H.Y., K.Y.; Supervision: K.Y.; Project Administration: K.Y., L.L.; Funding Acquisition: D.M.F.v.A., K.Y.

## Methods

### Protein purification and fluorescent labeling

A truncated *Cp*OGA (31-618 amino acids) and its derivative mutants were cloned into pET28 vector, with an N-terminal HaloTag and a C-terminal 6xHis-tag. The Rosetta (DE3) cells carrying the plasmids were grown overnight at 37 □ in 3 mL Luria-Bertani medium containing 50 μg/mL ampicillin (LB-Amp), and the overnight culture was reinoculated into a larger volume of fresh LB-Amp at 1 mL/L. Cells were then grown to an OD_600_ of 0.6, and the protein expression was induced with 0.5 mM of IPTG at 20 □ for 14 hours. Cells were harvested by centrifugation at 4000 g for 15 min at 4 □, and the pellets were resuspended at 10 mL/g in ice cold lysis buffer (20 mM Tris, 500 mM NaCl, 10 mM Imidazole at pH 8.0) supplemented with protease inhibitors (0.2 mM PMSF and 5 μM leupeptin). Cells were then lysed using high-pressure cell disrupter (Union-biotech) at 600 kpa for three times, and the lysate was cleared by centrifugation at 12000 g for 20 min at 4 □. The supernatant was collected and loaded onto 2 mL pre-equilibrated Ni-NTA agarose (Macherey-Nagel). The mixture was rotated at 4 □ for 2 h, and the agarose was washed with 5 mL wash buffer (200 mM Tris, 500 mM NaCl, 20 mM Imidazole at pH 7.9) for three times. The protein was eluted with 1 mL of elution buffer (20 mM Tris, 200 mM NaCl, 300 mM Imidazole at pH 7.9), and then dialysed into 40 mM Hepes (pH 7.4) and 150 mM KCl using dialysis tubing (SnakeSkin, Thermo Fisher). The HaloTag-*Cp*OGA proteins were mixed with equal molar of HaloTag Alexa Fluor 488 or 660 ligands (Promega) and incubated at room temperature for 30 min. The mixture was filtered through the G-50 column (GE Healthcare) at 2000 g for 2 min to remove the free dyes, aliquoted, flash frozen in liquid nitrogen, and stored at -80 □. The PCNA-mCherry protein was expressed and purified as previously reported ^36^.

### Cell lines

Hela cells (Meisen CTCC) were maintained as sub-confluent monolayers in Dulbecco’s modified Eagle’s medium (Gibco) with 10% fetal bovine serum (Biological Industries) at 37 □ with 5% CO_2_. The plasmids were transfected using Lipofectamine 2000 (Invitrogen).

### Fly stocks and genetic crosses

*Drosophila melanogaster* was raised on standard cornmeal-molasses-agar medium. The strains used in this study were as follow: *Sevelen* (used as the wild type in imaging), ;*Ogt*^H537A^;, MTD-gal4 (otu-gal4::VP16;gal4-nos;gal4::VP16-nos, BDSC, #31777), ;Da-gal4;, H2AvD-RFP (BDSC, #5905), uas-sh*sog* (BDSC, #67975), uas-GFP-Ph and uas-GFP-PC (kindly provided by Giacomo Cavalli), sogDistal>MS2 and sogIntronic>MS2 (kindly provided by Michael Levine), yw;MCP_Φ_NLS-mCherry (kindly provided by Hernan Garcia), ;sco/cyo;TM3/TM6B.

For the generation of transgenic lines, uas-*Cp*OGA^WT^, uas-*Cp*OGA^CD^, and uas-*Cp*OGA^DM^ sequences were codon optimized to *Drosophila melanogaster* using Jcat ^49^, and synthesized by Sangon Biotech Co., Ltd. HA and GFP tag were added to the N-terminal of *Cp*OGA sequences, and the fragments were cloned into pUASz attB vector (kindly provided by Hong Xu) using BamHI and XbaI. The constructs along with the helper plasmid were injected into y1,w67c23;P{CaryP}attP40 (UniHuaii). The resulted adult flies (G0) were crossed to double balancer to get the F1 generations.

For ChIP-seq and staining experiments, ;uas-*Cp*OGA,Da-gal4; lines were generated via recombination and the homozygous flies were used. For behavior assays, Da-gal4 virgins were mated with uas-*Cp*OGA^WT^ or uas-*Cp*OGA^DM^ males, and then the F1 was used in learning test. MTD-gal4 virgins were mated with uas-*Cp*OGA^WT^ or uas-*Cp*OGA^DM^ to generate the F1, and the F1 was mated to their siblings to generate the F2 for learning test. For MS2 live imaging, yw;MCPΦNLS-mCherry virgins were mated with homozygous males carrying MS2 alleles to image the nascent transcripts ^5^.

### Learning test

All behavior experiments were carried out at 25 °C and in an environmental chamber with 70% humidity as previously described ^41^. Each individual n consisted of approximately 200 flies (male:female=1:1), with half of the flies trained with one odor and half with the other odor. Flies were trained with a single training session and tested immediately. The odor avoidance to 3-octanol (OCT, no. 218405, Sigma-Aldrich) or 4-methylcyclohexanol (MCH, no. 153095, Sigma-Aldrich), and the shock avoidance to 60 V were quantified as previously described ^41^. To avoid experimental errors caused by the preference of flies to the odor or the environment, the results with OCT or MCH accompanying the electric shock were used alternately to calculate the performance index (PI). PI=[n(CS^-^)-n(CS^+^)]/[n(CS^+^)+n(CS^-^)]×100%. n(CS^+^) represents the number of flies choosing the odor coupled to the electric shock. n(CS^-^) represents the number of flies selecting the odor that was not coupled to the electric shock. The higher PI means stronger olfactory learning ability.

### Immunofluorescence

Hela cells were quickly rinsed with pre-warmed DPBS (Biological Industries), and then fixed with 10% formaldehyde in DPBS for 10 min. After three washes in DPBS, cells were permeabilized in 0.1% Triton X-100 for 10 min, and then washed with DPBS for three times before being blocked with 5% bovine serum albumin in DPBS for 30 min. The cells were incubated at 4 overnight with anti-O-GlcNAc antibody RL2 (1:400 ab2739, Abcam) or Alexa Fluor 660 labeled HaloTag-*Cp*OGA^CD^ (10 μg/mL). After three times wash with DPBS, the cells were either further incubated with fluorescently labeled secondary antibody for 1 h (1:500, Thermo Fisher) or directly incubated with DAPI (0.5 μg/mL) for 3 min to visualize the DNA. The cells were mounted with SlowFade Diamond antifade mountant (Thermo Fisher).

Embryos or polytene chromosome squash for immunostainings were prepared as previously described ^36^. The following primary antibodies were used: anti-phospho-Histone H3 (1:500, 9710s, CST), anti-H3K27me3 (1:100, C36B11, CST), anti-H2AK119ub (1:200, D27C4, CST). The Alexa Fluor 488 or Alexa Fluor 660 labelled HaloTag-*Cp*OGA^CD^ was used at 10 μg/mL.

### Microinjection and live imaging

The microinjection was performed as previously described ^36^. Briefly, embryos were manually dechorionated, aligned, taped onto a coverslip, desiccated for 4-6 min, and then embedded in halocarbon oil 27-halocarbon oil 700 (1:1, Sigma-Aldrich). The embryos were aged to cycle 9-10 before injection of fluorescent probes using a needle equipped on a manual micromanipulator (WPI) and controlled by the SYS-PV830 pneumatic pump (WPI). Alexa Fluor 488 or Alexa Fluor 660 labelled HaloTag-*Cp*OGA proteins were used at 10 μg/μL. The unlabelled *Cp*OGA^WT^ and *Cp*OGA^DM^ were used at 20 μg/μL. PCNA-mCherry was used at 2 μg/μL, and WGA (W21404, Thermo Fisher) was used at 1.25 μg/μL. The embryos were then imaged on ZEISS LSM880 confocal system with a 63x Plan-Apochromat 1.4 NA oil objective at room temperature. The GFP-Ph images were pseudo-colored with the green fire blue look-up table (LUT) in Fiji v.1.53f51.

### Western blot and Far Western

Embryos were collected on apple juice agar plates at 25 °C for 30 min and aged for 30 min, 90 min, 120 min, 180 min before being lysed in Lysis buffer (2% SDS, 10% glycerol, 62.5 mM Tris-HCl, pH 6.8) supplemented with 1x protease inhibitor cocktail (P8340, Sigma-Aldrich) and 50 μM Thiamet-G (s7213, Selleck) to detect the O-GlcNAcylation level. The protein concentrations were estimated using BCA protein assay (Beyotime), and 25 μg of the crude lysates were subjected to SDS-PAGE and transferred onto nitrocellulose membrane before immunoblotting with RL2 (1:1000, ab2739, Abcam) or CTD110.6 (1:1000, MABS1254, Sigma-Aldrich). Embryos expressing *Cp*OGA^WT^ or *Cp*OGA^DM^ were aged for 160 min before being lysed to perform western blot with anti-H3K27me3 (1:1000, C36B11, CST) or anti-H2AK119ub (1:1000, 8240S, CST). The secondary antibodies were used at 1:10000 (Thermo Fisher). The Far Western experiment was performed as previously described ^29^. The blot was incubated with 10 μg/mL pre-biotin-*Cp*OGA^CD^ for 1 h at room temperature followed by incubation with streptavidin-HRP (1:5000, M00091, GenScript) for 30 min.

### Embryo collection, embryo sorting, and chromatin preparation

Embryos were collected at 25°C for 30 min and aged for 160 min before being harvested and fixed. The embryos were then hand sorted in small batches under a light microscope (MZ61, Mshot) to remove those younger or older than the targeted age range based on morphology as previously described ^37^. After sorting, the embryos were stored at -80 °C. The chromatin for subsequent ChIP-seq experiment was prepared as described previously ^50^.

### ChIP-seq and data analysis

The chromatin was fragmented to sizes ranging from 100 to 300 bp using the Covaris S2 focus ultrasonicator (number of cycles=4, duty cycle=10%, intensity=5, ON=30 s, OFF=30 s). The ChIP was carried out as previously described ^50^ with 0.5 μg of anti-H3K27me3 (C36B11, CST), 2.7 μg of anti-H2AK119ub (D27C4, CST), or 5 μg of anti-pol II (ab817, Abcam). The sequencing libraries were prepared using the NEBNext Ultra II DNA Library Prep Kit for Illumina (NEB) following the manufacturer’s instructions. The libraries were sequenced on an Illumina Hiseq platform (Novagene, Tianjin, China).

The raw sequenced reads were cleaned using trim_galore, and then mapped to the reference *D. melanogaster* genome (dm6, version BDGP6) using Bowtie2 with the default parameters that look for multiple alignments but only report the one with the best mapping quality. Duplicate reads were removed using MarkDuplicates from gatk package v.4.1.4.1. Peaks were called using MACS (version 2.2.6) with parameters: -g dm –nomodel –broad –broad-cutoff 0.1. Bigwig tracks were generate using bamCompare from deeptools (parameters: --skipNAs –scaleFactorsMethod readCount –operation subtract). The ChIP-seq profiles were generated by computeMatrix and plotHeatmap in deepTools. IGV v.2.9.2 was used to visualize the bigwig tracks.

### Single embryo RNA-seq and RT-qPCR

Single embryo RNA-seq was performed as previously reported ^5^. RNA extraction was done with TRIzol (Invitrogen), and the total RNA was made into libraries for sequencing using the mRNA-seq Sample Preparation Kit (Illumina). The library was sequenced on an Illumina Hiseq platform (Novagene, Tianjin, China). The raw reads of RNA-seq were cleaned with trim_galore v0.6.0 with default parameters. The reads were then mapped to the dm6 genome assembly downloaded from UCSC with STAR v2.5.3a. Gene expression was quantified with featureCounts v1.6.5 using the dm6 RefSeq genes annotation file. Differential expression analysis was performed using DESeq2. Heatmap was created using R package ComplexHeatmap v2.10.0. GO analysis was performed using DAVID.

For RT-qPCR, RNA extracted by TRIzol was reverse transcribed to cDNA using the RevertAid First Strand cDNA Synthesis Kit (K1622, Thermo Fisher). The cDNA was then used as templates and qPCR was performed using the SYBR Green qPCR Master Mix (QST-100, SolomonBio) on the QuantStudio3 Real-Time PCR system (Applied Biosystems).

### Data and code availability

The sequencing data was deposited to the GEO database (accession number GSE197343). To review GEO accession GSE197343: go to https://www.ncbi.nlm.nih.gov/geo/query/acc.cgi?acc=GSE197343, and enter token: qhkpcsggpdqvdsh into the box.

## Supplemental information

Supplementary Table 1. Sequences of all the primers and protein used in this study.

Supplementary Table 2. Peaks of ChIP-seq, related to Fig 3D.

Supplementary Table 3. GO analysis and Annotation for Peaks of increased H3K27me3, related to Fig 3E-F.

Supplementary Table 4. Expression of the 651 genes with increased H3K27me3, related to Fig 3G.

Supplementary Table 5. Expression of the MBT genes with H3K27me3, related to Fig 3H and Fig S3I.

Supplementary Table 6. Expression of the total genes, related to Fig S3F.

Supplementary Table 7. Overlap between genes with inc_H3K27me3 and MBT active or MBT poised genes, related to Fig S3H.

Supplementary Movie 1. Learning test, related to Fig 4E.

---

**Figure S1.**
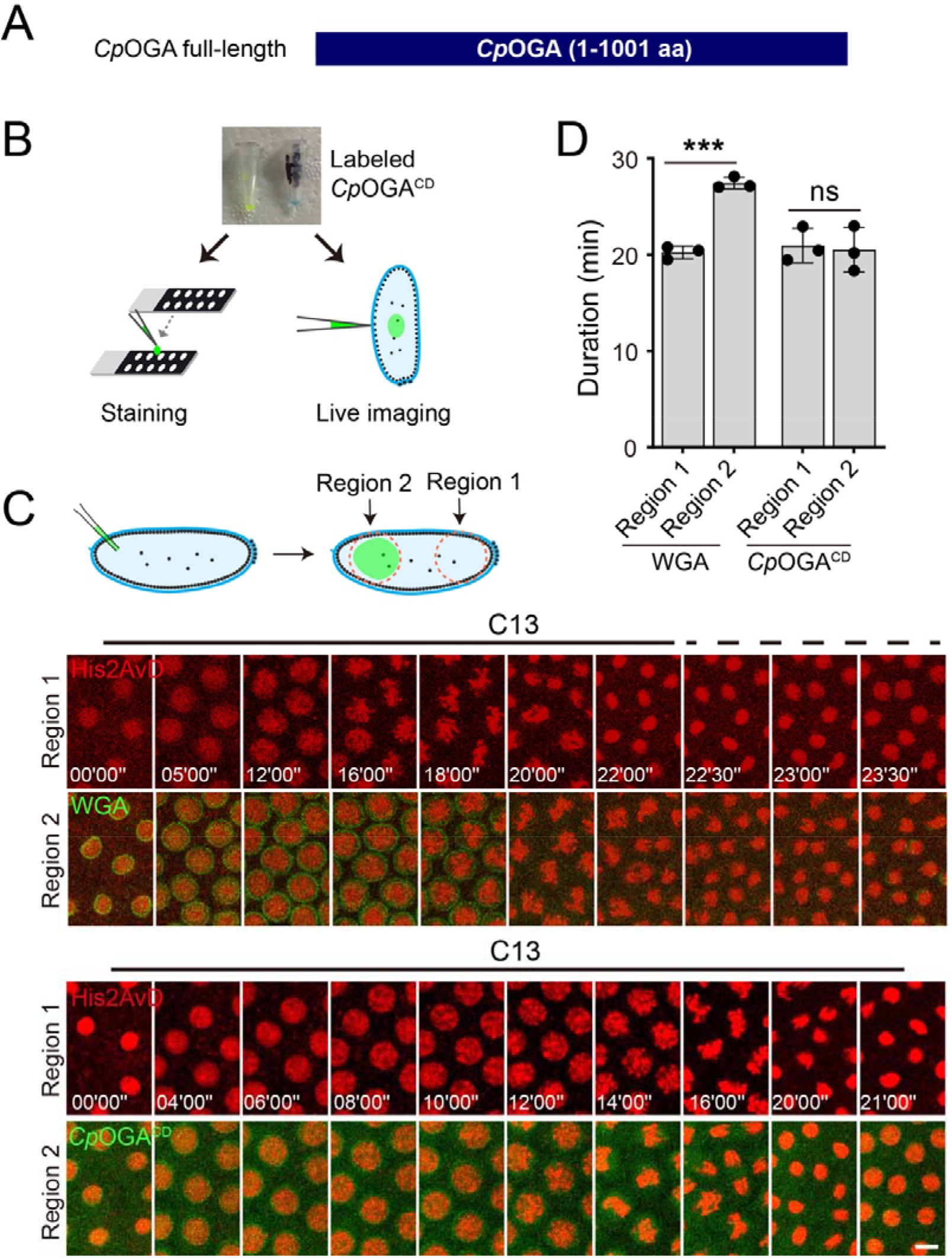
Live imaging of protein O-GlcNAcylation by fluorescently labeled *Cp*OGA^CD^ without disrupting the metasynchrony of embryonic cell divisions. **(A)** Schematic of the full-length *Cp*OGA (1-1001 aa). **(B)** Applications of the recombinant *Cp*OGA proteins in different experimental settings. **(C)** Characterization of the impact of WGA or *Cp*OGA^CD^ injection on the rapid and synchronous early embryonic cell cycles. WGA or *Cp*OGA^CD^ is injected at one end of the embryo during the interphase of cycle 12, and nuclei from both ends are imaged (region 1 and region 2). WGA or *Cp*OGA^CD^ is shown in green in the video frames, and His2AvD-RFP expressed from a transgene labeling the chromosomes in red. Relative time from the exit of mitosis 12 is given in minutes and seconds. Bar: 5 μm. **(D)** A compilation of timing results of the interphase of cycle 13 (C13). ns: not significant, \*\*\**p* < 0.001 by paired t-test. Error bars represent SD.

**Figure S2.**
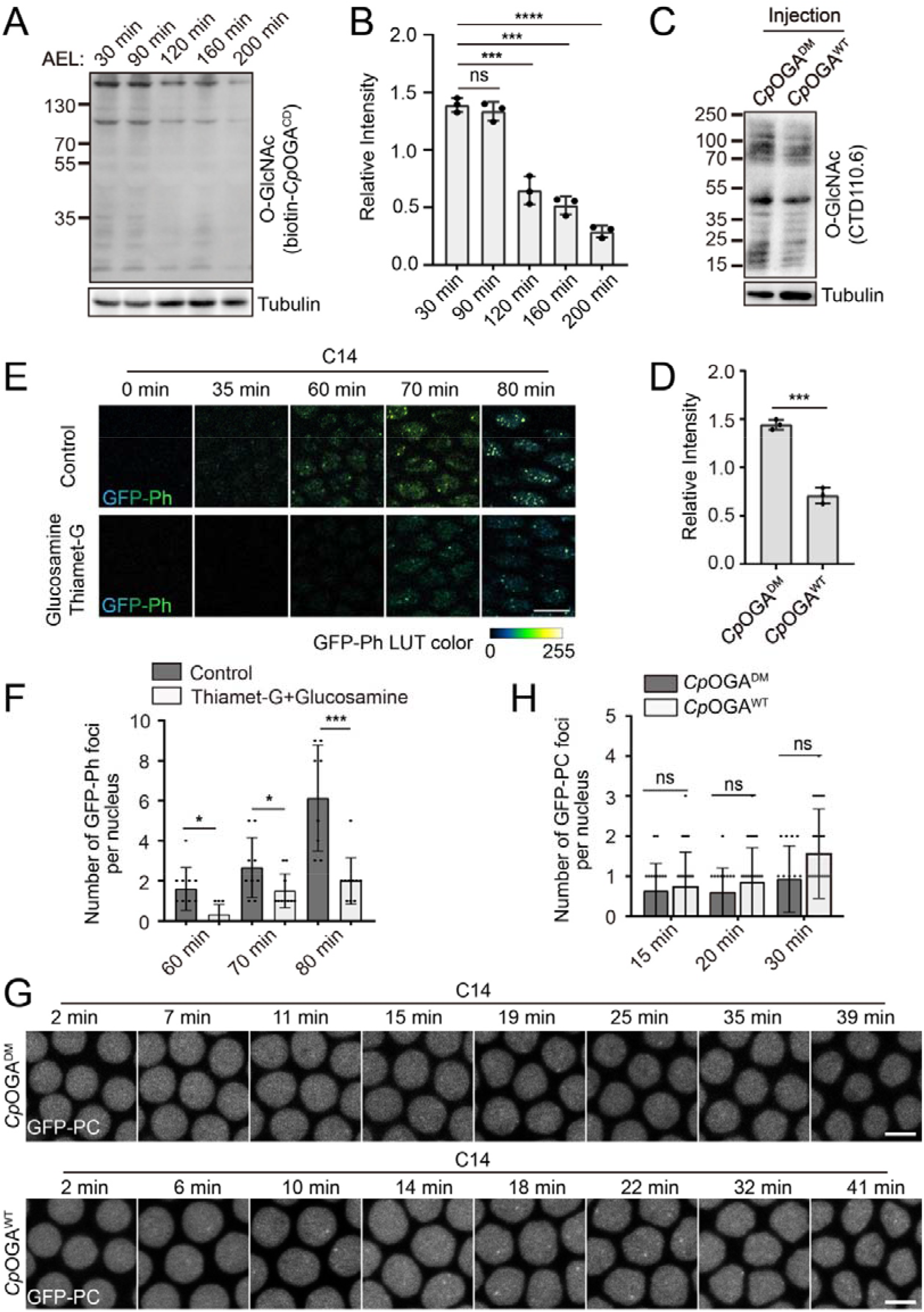
Altered kinetics of Ph bodies formation after manipulating the O-GlcNAcylation level. **(A)** Detection of total O-GlcNAcylation proteome at different timepoints after egg laying (AEL) by far western blot using biotin-*Cp*OGA^CD^. **(B)** Quantification of O-GlcNAcylated proteins relative to tubulin. **(C)** Western blot of O-GlcNAcylated proteins using anti-O-GlcNAc antibody CTD110.6 in embryos injected with *Cp*OGA^WT^ or control *Cp*OGA^DM^. **(D)** Quantification of O-GlcNAcylated proteins relative to tubulin. **(E)** Video frames of GFP-Ph from live embryos injected with control buffer or a mixture of Glucosamine and Thiamet-G. Relative times from the beginning of cycle 14 in minutes are given on the top. Bar: 10 μm. **(F)** Quantification of number of GFP-Ph foci per nucleus at the indicated time points. **(G)** Video frames from live imaging of GFP-PC in embryos injected with *Cp*OGA^WT^ or control *Cp*OGA^DM^. Elapsed times after entering cycle 14 are shown on the top. Bars: 5 μm. **(H)** Quantification of GFP-PC foci. ns: not significant,\**p* < 0.05, \*\**p* < 0.01, \*\*\**p* < 0.001, \*\*\*\**p* < 0.0001 by unpaired t-test. Error bars represent SD.

**Figure S3.**
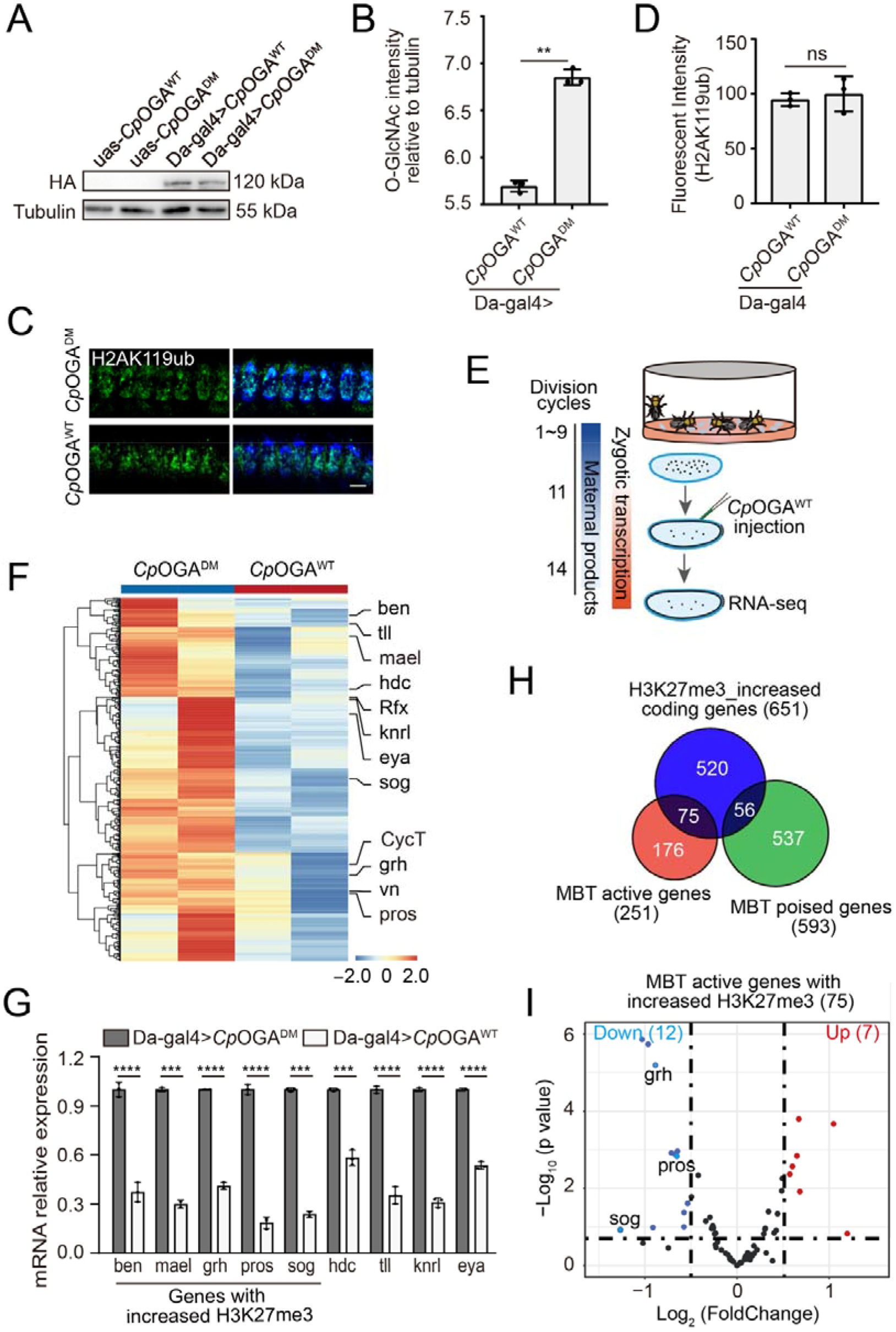
Changes in transcriptome after injection of exogenous *Cp*OGA. **(A)** Detection the Da-gal4 driven expression of *Cp*OGA^WT^ and *Cp*OGA^DM^ in *Drosophila* embryos by western blot with anti-HA antibody. **(B)** Quantification of western blots of O-GlcNAcylated protein relative to tubulin in embryos transgenically expressing *Cp*OGA^WT^ or control *Cp*OGA^DM^ with anti-O-GlcNAc antibody CTD110.6. **(C)** Immunostaining of embryos expressing *Cp*OGA^WT^ or *Cp*OGA^DM^ with anti-H2AK119ub antibody (green). Bar: 5 μm. **(D)** Quantification of the nuclear H2AK119ub fluorescent signals. **(E)** Schematic demonstrating the RNA-seq analysis of embryos injected with the indicated *Cp*OGA proteins. **(F)** Heatmap of the 535 downregulated genes in embryos injected with *Cp*OGA^WT^ versus control *Cp*OGA^DM^. See also Supplementary Table 6. **(G)** qPCR validation of a fraction of the downregulated genes with embryos transgenically expressing *Cp*OGA^WT^ or *Cp*OGA^DM^. **(H)** Analysis of overlap between the genes with increased H3K27me3 in *Cp*OGA^WT^ expressing embryos and the previously defined MBT active or MBT poised genes. See also Supplementary Table 7. **(I)** Volcano plot showing the differential expression of the 75 MBT active genes with increased H3K27me3. ns: not significant, \*\**p* < 0.01, \*\*\**p* < 0.001, \*\*\*\**p* < by unpaired t-test. Error bars represent SD.

**Figure S4.**
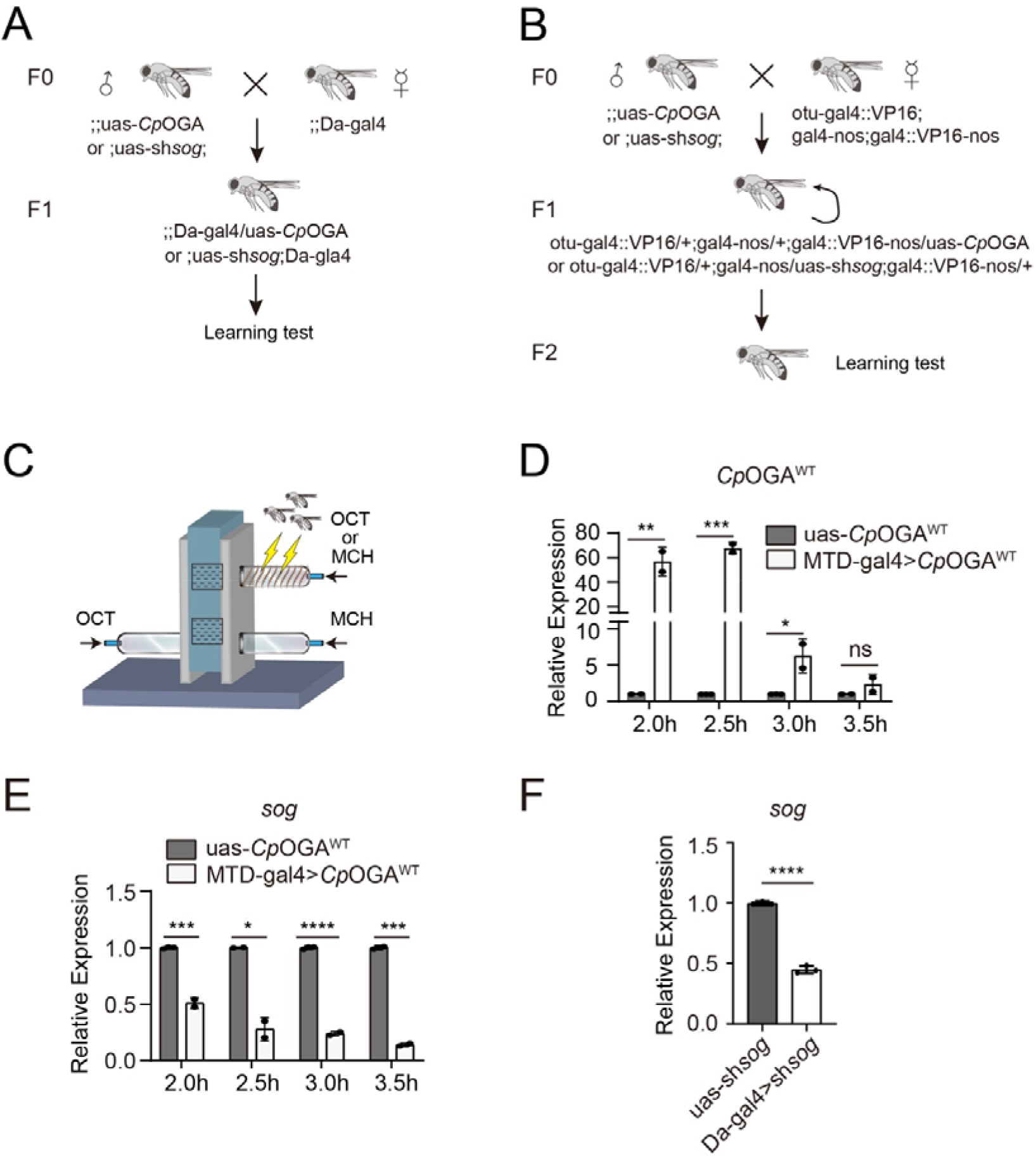
Modulation of *sog* transcription by protein O-GlcNAcylation. **(A-B)** Genetic cross schemes for flies used in the learning test. **(C)** Schematic of the device for *Drosophila* learning test. OCT: 3-octanol. MCH: 4-methylcyclohexanol. **(D)** qPCR analysis of *Cp*OGA^WT^ expression at the indicated timepoints after egg laying. **(E)** qPCR analysis of *sog* expression in embryos collected from the indicated genotypes at different timepoints. **(F)** qPCR analysis of *sog* expression after shRNA mediated knockdown. ns: not significant, \**p* < 0.05, \*\**p* < 0.01, \*\*\**p* < 0.001, \*\*\*\**p* < 0.0001 by unpaired t-test. Error bars represent SD.

## References

1. O’Farrell, P.H. (2015). Growing an Embryo from a Single Cell: A Hurdle in Animal Life. Cold Spring Harb Perspect Biol 7. 10.1101/cshperspect.a019042.

2. Farrell, J.A., and O’Farrell, P.H. (2014). From egg to gastrula: how the cell cycle is remodeled during the Drosophila mid-blastula transition. Annual review of genetics 48, 269–294. 10.1146/annurev-genet-111212-133531.

3. Yuan, K., Seller, C.A., Shermoen, A.W., and O’Farrell, P.H. (2016). Timing the Drosophila Mid-Blastula Transition: A Cell Cycle-Centered View. Trends Genet 32, 496–507. 10.1016/j.tig.2016.05.006.

4. Collart, C., Allen, G.E., Bradshaw, C.R., Smith, J.C., and Zegerman, P. (2013). Titration of four replication factors is essential for the Xenopus laevis midblastula transition. Science 341, 893–896. 10.1126/science.1241530.

5. Strong, I.J.T., Lei, X., Chen, F., Yuan, K., and O’Farrell, P.H. (2020). Interphase-arrested Drosophila embryos activate zygotic gene expression and initiate mid-blastula transition events at a low nuclear-cytoplasmic ratio. PLoS Biol 18, e3000891. 10.1371/journal.pbio.3000891.

6. Shindo, Y., and Amodeo, A.A. (2021). Excess histone H3 is a competitive Chk1 inhibitor that controls cell-cycle remodeling in the early Drosophila embryo. Current biology : CB 31, 2633–2642 e2636. 10.1016/j.cub.2021.03.035.

7. Djabrayan, N.J., Smits, C.M., Krajnc, M., Stern, T., Yamada, S., Lemon, W.C., Keller, P.J., Rushlow, C.A., and Shvartsman, S.Y. (2019). Metabolic Regulation of Developmental Cell Cycles and Zygotic Transcription. Current biology : CB 29, 1193–1198 e1195. 10.1016/j.cub.2019.02.028.

8. Liu, B., Winkler, F., Herde, M., Witte, C.P., and Grosshans, J. (2019). A Link between Deoxyribonucleotide Metabolites and Embryonic Cell-Cycle Control. Current biology : CB 29, 1187–1192 e1183. 10.1016/j.cub.2019.02.021.

9. Vocadlo, D.J. (2012). O-GlcNAc processing enzymes: catalytic mechanisms, substrate specificity, and enzyme regulation. Curr Opin Chem Biol 16, 488–497. 10.1016/j.cbpa.2012.10.021.

10. Hart, G.W. (2014). Three Decades of Research on O-GlcNAcylation - A Major Nutrient Sensor That Regulates Signaling, Transcription and Cellular Metabolism. Front Endocrinol (Lausanne) 5, 183. 10.3389/fendo.2014.00183.

11. Olivier-Van Stichelen, S., and Hanover, J.A. (2015). You are what you eat: O-linked N-acetylglucosamine in disease, development and epigenetics. Curr Opin Clin Nutr Metab Care 18, 339–345. 10.1097/MCO.0000000000000188.

12. Hardiville, S., and Hart, G.W. (2016). Nutrient regulation of gene expression by O-GlcNAcylation of chromatin. Curr Opin Chem Biol 33, 88–94. 10.1016/j.cbpa.2016.06.005.

13. Shafi, R., Iyer, S.P., Ellies, L.G., O’Donnell, N., Marek, K.W., Chui, D., Hart, G.W., and Marth, J.D. (2000). The O-GlcNAc transferase gene resides on the X chromosome and is essential for embryonic stem cell viability and mouse ontogeny. Proc Natl Acad Sci U S A 97, 5735–5739. 10.1073/pnas.100471497.

14. Vaidyanathan, K., Niranjan, T., Selvan, N., Teo, C.F., May, M., Patel, S., Weatherly, B., Skinner, C., Opitz, J., Carey, J., et al. (2017). Identification and characterization of a missense mutation in the O-linked beta-N-acetylglucosamine (O-GlcNAc) transferase gene that segregates with X-linked intellectual disability. J Biol Chem 292, 8948–8963. 10.1074/jbc.M116.771030.

15. Willems, A.P., Gundogdu, M., Kempers, M.J.E., Giltay, J.C., Pfundt, R., Elferink, M., Loza, B.F., Fuijkschot, J., Ferenbach, A.T., van Gassen, K.L.I., et al. (2017). Mutations in N-acetylglucosamine (O-GlcNAc) transferase in patients with X-linked intellectual disability. J Biol Chem 292, 12621–12631. 10.1074/jbc.M117.790097.

16. Pravata, V.M., Muha, V., Gundogdu, M., Ferenbach, A.T., Kakade, P.S., Vandadi, V., Wilmes, A.C., Borodkin, V.S., Joss, S., Stavridis, M.P., and van Aalten, D.M.F. (2019). Catalytic deficiency of O-GlcNAc transferase leads to X-linked intellectual disability. Proc Natl Acad Sci U S A 116, 14961–14970. 10.1073/pnas.1900065116.

17. Pravata, V.M., Gundogdu, M., Bartual, S.G., Ferenbach, A.T., Stavridis, M., Ounap, K., Pajusalu, S., Zordania, R., Wojcik, M.H., and van Aalten, D.M.F. (2020). A missense mutation in the catalytic domain of O-GlcNAc transferase links perturbations in protein O-GlcNAcylation to X-linked intellectual disability. FEBS Lett 594, 717–727. 10.1002/1873-3468.13640.

18. Pravata, V.M., Omelkova, M., Stavridis, M.P., Desbiens, C.M., Stephen, H.M., Lefeber, D.J., Gecz, J., Gundogdu, M., Ounap, K., Joss, S., et al. (2020). An intellectual disability syndrome with single-nucleotide variants in O-GlcNAc transferase. Eur J Hum Genet. 10.1038/s41431-020-0589-9.

19. Ingham, P.W. (1984). A gene that regulates the bithorax complex differentially in larval and adult cells of Drosophila. Cell 37, 815–823.

20. Sinclair, D.A., Syrzycka, M., Macauley, M.S., Rastgardani, T., Komljenovic, I., Vocadlo, D.J., Brock, H.W., and Honda, B.M. (2009). Drosophila O-GlcNAc transferase (OGT) is encoded by the Polycomb group (PcG) gene, super sex combs (sxc). Proc Natl Acad Sci U S A 106, 13427–13432. 10.1073/pnas.0904638106.

21. Gambetta, M.C., Oktaba, K., and Muller, J. (2009). Essential role of the glycosyltransferase sxc/Ogt in polycomb repression. Science 325, 93–96. 10.1126/science.1169727.

22. Gambetta, M.C., and Muller, J. (2014). O-GlcNAcylation prevents aggregation of the Polycomb group repressor polyhomeotic. Dev Cell 31, 629–639. 10.1016/j.devcel.2014.10.020.

23. Ma, J., and Hart, G.W. (2014). O-GlcNAc profiling: from proteins to proteomes. Clin Proteomics 11, 8. 10.1186/1559-0275-11-8.

24. Lin, W., Gao, L., and Chen, X. (2015). Protein-Specific Imaging of O-GlcNAcylation in Single Cells. Chembiochem 16, 2571–2575. 10.1002/cbic.201500544.

25. Wu, Z.L., Tatge, T.J., Grill, A.E., and Zou, Y. (2018). Detecting and Imaging O-GlcNAc Sites Using Glycosyltransferases: A Systematic Approach to Study O-GlcNAc. Cell Chem Biol 25, 1428–1435 e1423. 10.1016/j.chembiol.2018.07.007.

26. Selvan, N., Williamson, R., Mariappa, D., Campbell, D.G., Gourlay, R., Ferenbach, A.T., Aristotelous, T., Hopkins-Navratilova, I., Trost, M., and van Aalten, D.M.F. (2017). A mutant O-GlcNAcase enriches Drosophila developmental regulators. Nat Chem Biol 13, 882–887. 10.1038/nchembio.2404.

27. Tan, H.Y., Eskandari, R., Shen, D., Zhu, Y., Liu, T.W., Willems, L.I., Alteen, M.G., Madden, Z., and Vocadlo, D.J. (2018). Direct One-Step Fluorescent Labeling of O-GlcNAc-Modified Proteins in Live Cells Using Metabolic Intermediates. J Am Chem Soc 140, 15300–15308. 10.1021/jacs.8b08260.

28. Rao, F.V., Dorfmueller, H.C., Villa, F., Allwood, M., Eggleston, I.M., and van Aalten, D.M. (2006). Structural insights into the mechanism and inhibition of eukaryotic O-GlcNAc hydrolysis. EMBO J 25, 1569–1578. 10.1038/sj.emboj.7601026.

29. Mariappa, D., Selvan, N., Borodkin, V., Alonso, J., Ferenbach, A.T., Shepherd, C., Navratilova, I.H., and vanAalten, D.M.F. (2015). A mutant O-GlcNAcase as a probe to reveal global dynamics of protein O-GlcNAcylation during Drosophila embryonic development. Biochem J 470, 255–262. 10.1042/BJ20150610.

30. Fu, C., and van Aalten, D.M.F. (2020). Native detection of protein O-GlcNAcylation by gel electrophoresis. Analyst 145, 6826–6830. 10.1039/c9an02506e.

31. Mariappa, D., Zheng, X., Schimpl, M., Raimi, O., Ferenbach, A.T., Muller, H.A., and van Aalten, D.M. (2015). Dual functionality of O-GlcNAc transferase is required for Drosophila development. Open Biol 5, 150234. 10.1098/rsob.150234.

32. Yuan, K., and O’Farrell, P.H. (2015). Cyclin B3 Is a Mitotic Cyclin that Promotes the Metaphase-Anaphase Transition. Current biology : CB 25, 811–816. 10.1016/j.cub.2015.01.053.

33. Deneke, V.E., Melbinger, A., Vergassola, M., and Di Talia, S. (2016). Waves of Cdk1 Activity in S Phase Synchronize the Cell Cycle in Drosophila Embryos. Dev Cell 38, 399–412. 10.1016/j.devcel.2016.07.023.

34. Deneke, V.E., Puliafito, A., Krueger, D., Narla, A.V., De Simone, A., Primo, L., Vergassola, M., De Renzis, S., and Di Talia, S. (2019). Self-Organized Nuclear Positioning Synchronizes the Cell Cycle in Drosophila Embryos. Cell 177, 925–941 e917. 10.1016/j.cell.2019.03.007.

35. Cheutin, T., and Cavalli, G. (2012). Progressive polycomb assembly on H3K27me3 compartments generates polycomb bodies with developmentally regulated motion. PLoS Genet 8, e1002465. 10.1371/journal.pgen.1002465.

36. Yuan, K., and O’Farrell, P.H. (2016). TALE-light imaging reveals maternally guided, H3K9me2/3-independent emergence of functional heterochromatin in Drosophila embryos. Genes Dev 30, 579–593. 10.1101/gad.272237.115.

37. Chen, K., Johnston, J., Shao, W., Meier, S., Staber, C., and Zeitlinger, J. (2013). A global change in RNA polymerase II pausing during the Drosophila midblastula transition. Elife 2, e00861. 10.7554/eLife.00861.

38. Gaertner, B., and Zeitlinger, J. (2014). RNA polymerase II pausing during development. Development 141, 1179–1183. 10.1242/dev.088492.

39. Holley, S.A., Jackson, P.D., Sasai, Y., Lu, B., De Robertis, E.M., Hoffmann, F.M., and Ferguson, E.L. (1995). A conserved system for dorsal-ventral patterning in insects and vertebrates involving sog and chordin. Nature 376, 249–253. 10.1038/376249a0.

40. Biehs, B., Francois, V., and Bier, E. (1996). The Drosophila short gastrulation gene prevents Dpp from autoactivating and suppressing neurogenesis in the neuroectoderm. Genes Dev 10, 2922–2934. 10.1101/gad.10.22.2922.

41. Jia, J., He, L., Yang, J., Shuai, Y., Yang, J., Wu, Y., Liu, X., Chen, T., Wang, G., Wang, X., et al. (2021). A pair of dopamine neurons mediate chronic stress signals to induce learning deficit in Drosophila melanogaster. Proc Natl Acad Sci U S A 118. 10.1073/pnas.2023674118.

42. Schulz, K.N., and Harrison, M.M. (2019). Mechanisms regulating zygotic genome activation. Nat Rev Genet 20, 221–234. 10.1038/s41576-018-0087-x.

43. Kim, C.A., Gingery, M., Pilpa, R.M., and Bowie, J.U. (2002). The SAM domain of polyhomeotic forms a helical polymer. Nat Struct Biol 9, 453–457. 10.1038/nsb802.

44. Humphreys, G.B., Jud, M.C., Monroe, K.M., Kimball, S.S., Higley, M., Shipley, D., Vrablik, M.C., Bates, K.L., and Letsou, A. (2013). Mummy, A UDP-N-acetylglucosamine pyrophosphorylase, modulates DPP signaling in the embryonic epidermis of Drosophila. Dev Biol 381, 434–445. 10.1016/j.ydbio.2013.06.006.

45. Moulton, M.J., Humphreys, G.B., Kim, A., and Letsou, A. (2020). O-GlcNAcylation Dampens Dpp/BMP Signaling to Ensure Proper Drosophila Embryonic Development. Dev Cell 53, 330–343 e333. 10.1016/j.devcel.2020.04.001.

46. Galeone, A., Han, S.Y., Huang, C., Hosomi, A., Suzuki, T., and Jafar-Nejad, H. (2017). Tissue-specific regulation of BMP signaling by Drosophila N-glycanase 1. Elife 6. 10.7554/eLife.27612.

47. Negreiros, E., Herszterg, S., Kang, K.H., Camara, A., Dias, W.B., Carneiro, K., Bier, E., Todeschini, A.R., and Araujo, H. (2018). N-linked glycosylation restricts the function of Short gastrulation to bind and shuttle BMPs. Development 145. 10.1242/dev.167338.

48. Galeone, A., Adams, J.M., Matsuda, S., Presa, M.F., Pandey, A., Han, S.Y., Tachida, Y., Hirayama, H., Vaccari, T., Suzuki, T., et al. (2020). Regulation of BMP4/Dpp retrotranslocation and signaling by deglycosylation. Elife 9. 10.7554/eLife.55596.

49. Grote, A., Hiller, K., Scheer, M., Munch, R., Nortemann, B., Hempel, D.C., and Jahn, D. (2005). JCat: a novel tool to adapt codon usage of a target gene to its potential expression host. Nucleic Acids Res 33, W526–531. 10.1093/nar/gki376.

50. Loubiere, V., Delest, A., Schuettengruber, B., Martinez, A.M., and Cavalli, G. (2017). Chromatin Immunoprecipitation Experiments from Whole Drosophila Embryos or Larval Imaginal Discs. Bio Protoc 7, e2327. 10.21769/BioProtoc.2327.

